# The application of zeta diversity as a continuous measure of compositional change in ecology

**DOI:** 10.1101/216580

**Authors:** Melodie A. Mcgeoch, Guillaume Latombe, Nigel R. Andrew, Shinichi Nakagawa, David A. Nipperess, Mariona Roige, Ezequiel M. Marzinelli, Alexandra H. Campbell, Adriana Vergés, Torsten Thomas, Peter D. Steinberg, Katherine E. Selwood, Cang Hui

## Abstract

Zeta diversity provides the average number of shared species across *n* sites (or shared operational taxonomic units (OTUs) across *n* cases). It quantifies the variation in species composition of multiple assemblages in space and time to capture the contribution of the full suite of narrow, intermediate and wide-ranging species to biotic heterogeneity. Zeta diversity was proposed for measuring compositional turnover in plant and animal assemblages, but is equally relevant for application to any biological system that can be characterised by a row by column incidence matrix. Here we illustrate the application of zeta diversity to explore compositional change in empirical data, and how observed patterns may be interpreted. We use 10 datasets from a broad range of scales and levels of biological organisation – from DNA molecules to microbes, plants and birds – including one of the original data sets used by R.H. Whittaker in the 1960’s to express compositional change and distance decay using beta diversity. The applications show (i) how different sampling schemes used during the calculation of zeta diversity may be appropriate for different data types and ecological questions, (ii) how higher orders of zeta may in some cases better detect shifts, transitions or periodicity, and importantly (iii) the relative roles of rare versus common species in driving patterns of compositional change. By exploring the application of zeta diversity across this broad range of contexts, our goal is to demonstrate its value as a tool for understanding continuous biodiversity turnover and as a metric for filling the empirical gap that exists on spatial or temporal change in compositional diversity.

## INTRODUCTION

Changes in the composition of diversity in space and time, along with richness, abundance and biomass, are critical to understanding what drives biodiversity and the ways that humans are transforming it (McGill et al. 2015). Interest in measuring and understanding the way in which species composition changes in space and time has risen exponentially over the last two decades (Anderson et al. 2011, Shimadzu et al. 2015, Myers and LaManna 2016, Socolar et al. 2016). Compositional change is not only relevant to species diversity, but to other levels of biological organisation, including molecular, genetic and phylogenetic diversity (e.g. Nipperess et al. 2012, Thomas et al. 2016), as well as social phenomena such as cultural diversity, economic development, collaboration and societal instability (e.g. Nettle et al. 2007, Vaz et al. 2017). The concept of turnover^1^ in the identity of elements is therefore relevant across a broad range of biological and socioecological systems that span multiple scales (Arita et al. 2012).

Zeta diversity was recently proposed as a concept that focusses attention on multi-site, cross-scale, assemblage patterns of turnover in biodiversity, with the purpose of better understanding how biodiversity is structured (Hui and McGeoch 2014). The zeta diversity measure quantifies the number of species shared by any given number of sites, and calculates all possible components from assemblage partitioning. Compositional, or incidence-based, turnover has traditionally been measured using metrics based on pairwise comparisons (*i*=2) of species incidence across sites or samples (Jost et al. 2010), commonly referred to as beta diversity (e.g. Jaccard dissimilarity). Differences in species composition between pairs of sites are driven largely by rare species rather than common ones (which are, by definition, shared by large numbers of sites). Comparisons of compositional change across *i* > 2 sites thus provides information on the contribution of increasingly more common (widespread) species in the assemblage to turnover.

The mathematical necessity of multiple site turnover measures, such as zeta diversity, has been shown. With information on only the alpha and all pairwise beta components in a community, it is not possible to know the full complement of partitions across multiple sites (Hui and McGeoch 2014). Dissimilarity indices based on combinations of multiple sites have been proposed (e.g. Diserud and Ødegaard 2007, Baselga et al. 2007, 2013), but provide a single measure of compositional turnover for a set of sites. By contrast, zeta diversity as a concept for the first time draws attention to the value of the full suite of multisite comparisons to quantifying compositional change. By incrementally increasing the number of sites and quantifying compositional change at each step, zeta diversity provides information on the full spectrum of rare to intermediate and common species as they contribute to driving compositional change. As such it provides a more comprehensive picture of turnover than a single aggregated value for compositional comparison. As a measure, zeta diversity (*ζ*_*i*_) enables this exploration of how incidence-based composition changes with both scale and number of sites *i* involved (Hui and McGeoch 2014).

The applied value of zeta diversity has to date also been shown in particular cases, for example as a measure of similarity and uncertainty in pest profile analysis (Roige et al. 2017), to measure field-specific interdisciplinarity (Vaz et al. 2017) and to upscale estimates of biodiversity (Kunin et al. in press). However, the main applications of zeta diversity (zeta decline and zeta decay) to classic incidence matrices in ecology, and how these are interpreted, has not yet been systematically illustrated. Using a range of levels of biological organisation, we show how zeta diversity can be applied and interpreted to provide insights on the nature of biotic heterogeneity. Building on Hui and McGeoch (2014), we also introduce for the first time the species retention rate using the zeta ratio, which quantifies relative rate of turnover in rare and common species. Zeta diversity is one among several developments in the field (e.g. Baselga 2010, 2013). While recognizing these developments, the aim here is not to contrast them, but rather to enable ecologists to further explore the structurally novel value and ecological insights provided by zeta diversity (Appendix S1 provides an illustrative approach).

## CALCULATING ZETA DIVERSITY

### Analysis

Throughout we use ‘OTU’ (operational taxonomic unit) to refer to species or other levels of biological organisation, ‘case’ to refer to site, sample, assemblage or other unit of comparison, and ‘community’ to refer to the OTU by case matrix. Zeta diversity (*ζ*_i_) is the mean number of OTUs shared by *i* number of cases, with *i* referred to as the zeta order, *ζ*_1_ (where *i* = 1) is the mean number of OTUs across all cases (or alpha diversity). The first-order of zeta diversity (*ζ*_1_), or average species richness, is thus equivalent to alpha, and the total observed or estimated richness across all sites or assemblages, as usual, represents gamma diversity. Incidence-based, pairwise beta similarity metrics are equivalent to *ζ*_2_ (Hui and McGeoch 2014), and higher orders of zeta (*i* > 2) represent the contribution of increasingly widespread (common) OTUs to compositional change. Analyses can be performed either using raw zeta, i.e. the absolute number of OTUs shared by cases, or on transformations of zeta.

Richness can vary substantially across sites and assemblages and, if desired, normalised zeta provides one option for dealing with richness difference effects (see for example Roige et al. 2017, depending on the study objective, other approaches are possible, e.g. Latombe et al. 2017). Normalised zeta is ζ_ij_/γ_j_, where ζ_ij_ is the number of species shared by the *i* sites in the specific combination *j*, and where γ_j_ (gamma diversity) is the total number of OTUs over the cases in the specific combination *j* (i.e. the gamma diversity of the combination). Normalised zeta is useful for comparing communities with large differences in richness, or where richness-independent patterns of turnover are of interest. The number of orders included in calculation of zeta is decided based on the dataset and question of interest, and at a maximum will be the total number of cases. If zeta reaches zero after *i* orders, i.e. no OTU is shared by more than *i* cases, there is of course no information to be gained by expressing it for orders beyond this.

All analyses were performed using the *zetadiv* package V.1.0 (Latombe et al. 2016), in R (R Core Team 2013). For each dataset, only those results that best illustrate each of the particular zeta diversity applications are discussed (full results in Supplementary Information). Like alpha and beta diversity, zeta diversity can be used in a wide variety of analyses, to quantify multiple facets of biodiversity. The two main applications are explored in detail in this paper, (1) zeta decline, including the OTU retention rate based on the zeta ratio, and (2) zeta decay over space or time.

### Data structure and sub-sampling schemes

For any dataset, the combination of a specific data structure and choice of sub-sampling scheme results in different possible pathways for expressing zeta diversity (Fig. 1, Appendix S1). The sub-sampling scheme for *i* cases (Fig. 1) has a significant effect on the value and interpretation of diversity patterns (Scheiner et al. 2011), including those quantified using zeta diversity. The data sub-sampling scheme may encompass all (or a random selection) or only a subset of possible combinations of *i* samples, and partially depend on the spatial or temporal structure of the data (Fig. 1). The main data sub-sampling schemes are all combinations (ALL), nearest neighbours (NON - non-directional or DIR - directional), and fixed point origin (FPO) or fixed edge origin (FEO) (Fig. 1). When zeta decline is calculated using the ALL combinations scheme (Fig. 1a, g), it provides an average expectation of compositional change in the data and could be considered as the lower bound (least shared OTUs) of expected turnover against structured sample designs. In cases where sites or surveys are positioned across a spatial or temporal gradient, and zeta is calculated using nearest neighbours (DIR or NON schemes, Figure 1b,c,e), zeta diversity will decline at a comparatively slower rate. This is due to the constraints imposed by this spatial or temporal dependence in the data sub-sampling scheme (versus the ALL scheme that considers combinations of sites that may be far from each other, and are therefore less likely to share OTUs than close sites). Zeta decline (using the ALL sub-sampling scheme) can thus be considered a null model against which scale or environmental mechanisms hypothesised to be responsible for driving patterns of turnover can be tested. Other sub-sampling schemes may be envisaged for more specific applications.

**FIG. 1.**
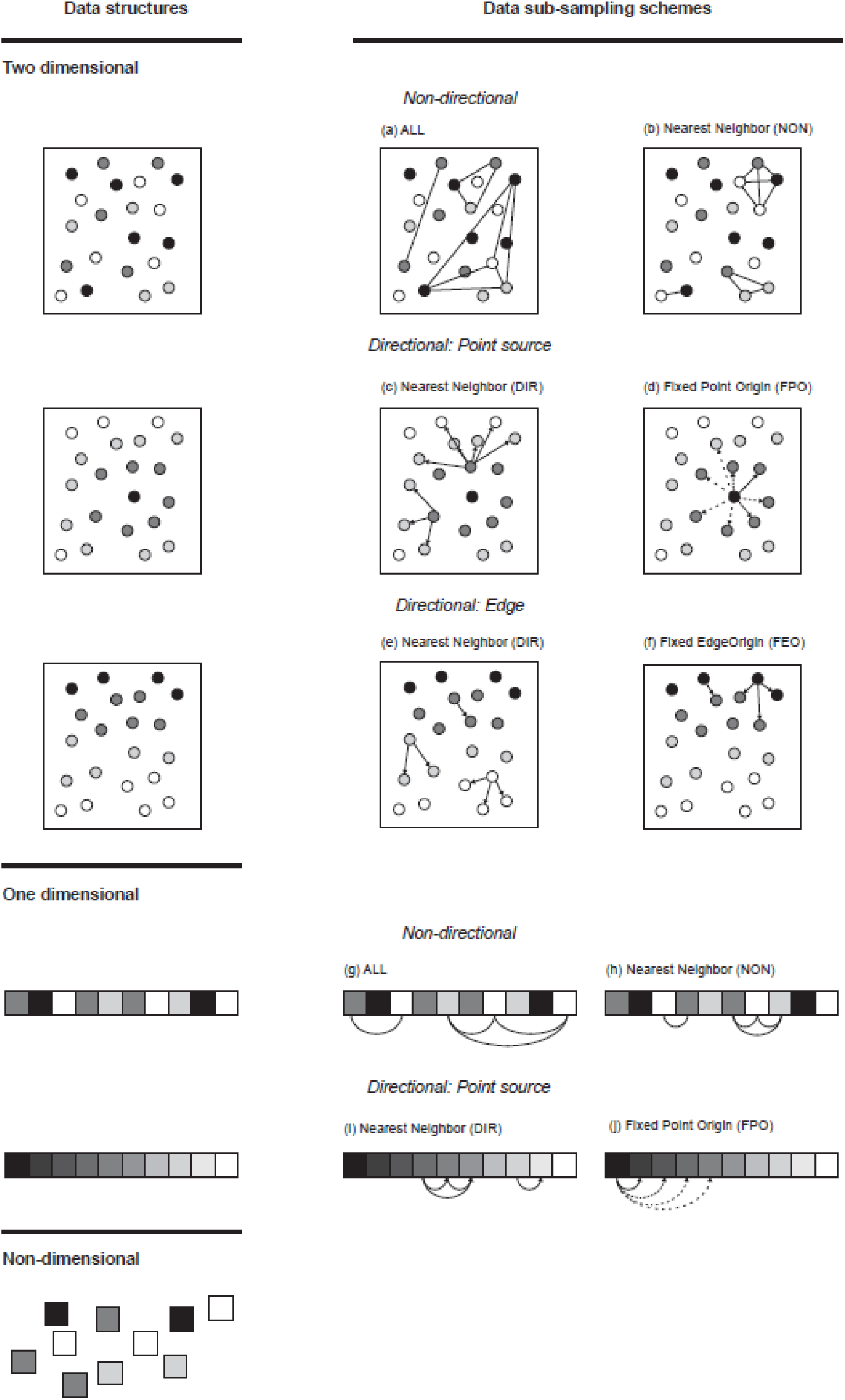
One- and two- dimensional data structures and alternative data sub-sampling schemes for estimating compositional turnover using zeta diversity (jointly referred to as the pathway for expressing zeta diversity). Data may include broad geographic regions encompassing spatially homogenous or heterogeneous environments (which may include multiple complex gradients as in (a,b)), independent units hosting a community (e.g. islands, hosts of parasite or bacterial communities or genomes) or linear habitats (e.g. coastlines or ecotones (a,b). The lines between sites are not comprehensive and simply show how sites may be combined for the calculation of zeta diversity. Directional structures are those where there are known or designed directional gradients of interest (c-f,i,j), e.g. a one or two dimensional change in environmental condition away from a point source (d), gradients perpendicular to an edge or ecotone (f), or a time series or transect along an environmental gradient (i,j). Non-dimensional schemes are those where no, or no single, environmental or spatial gradient is of concern or interest (sample units may also be discrete with their relative spatial position of no interest).

The choice of the data pathway, i.e. the combination of data structure and data sub- sampling scheme, will affect the outcome and is therefore important to consider *a priori* to ensure selection of the most appropriate pathway for the data and hypothesis of interest.

### Data

Ten datasets were used to demonstrate the application of zeta diversity and represent a range of taxa, levels of biological organisation and spatial or temporal scales (Table 1). The data sets also encompass a broad range of OTU richness (39 to 1804) and numbers of cases (< 20 to >1000). Each data set was structured as an OTU by case matrix with non-zero marginal totals. Singletons (OTUs present at only a single site) were removed from some datasets, especially where they are likely to be a result of under sampling or sampling bias (for further detail on treatment of individual datasets see Appendix S2). While each of these datasets described below potentially warrants a dedicated examination of compositional turnover of its own, here we use the diversity of cases and data structures to illustrate the application, and interpretation of zeta diversity and not to test data set-specific hypotheses per se.

**TABLE 1.**
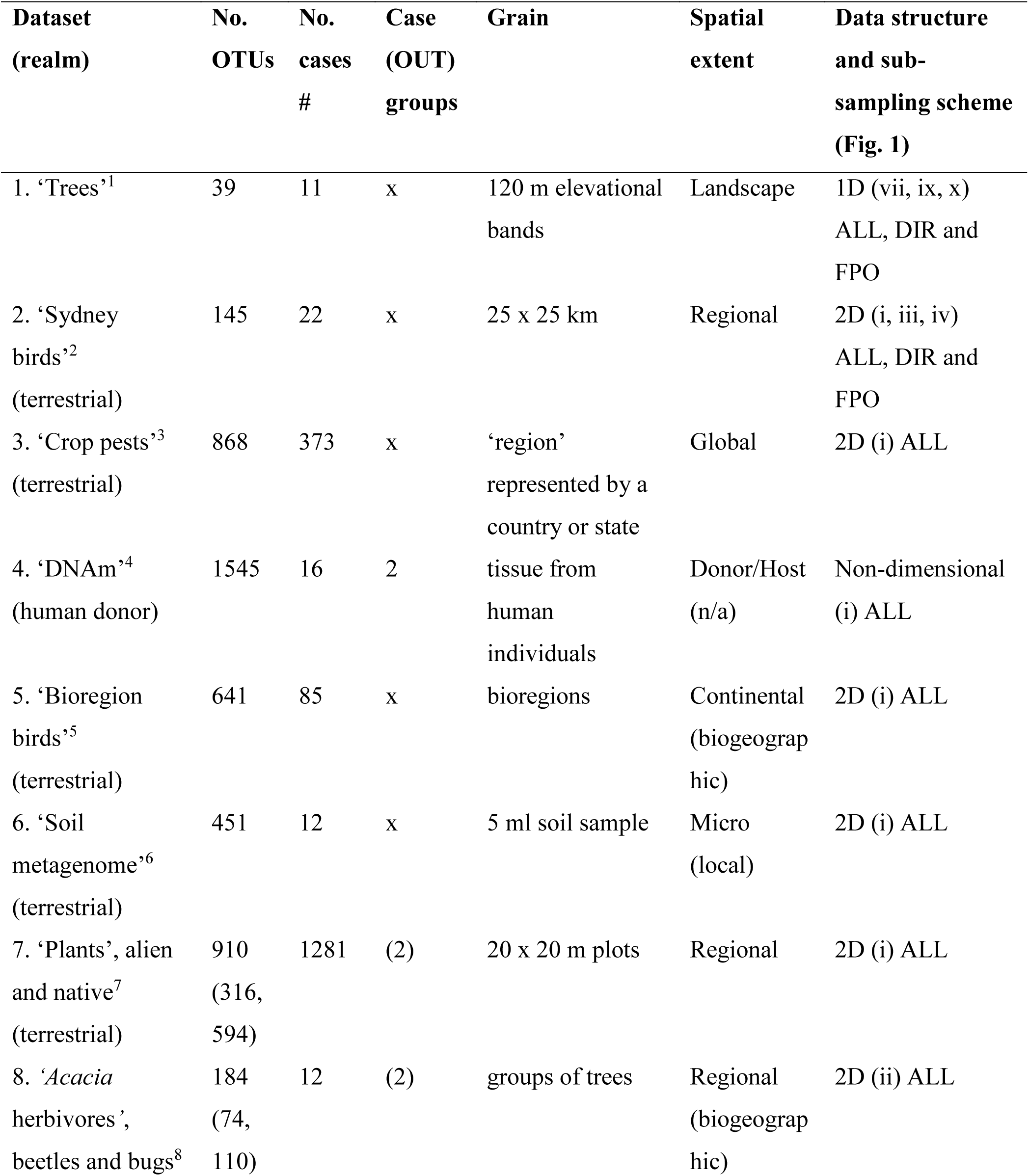
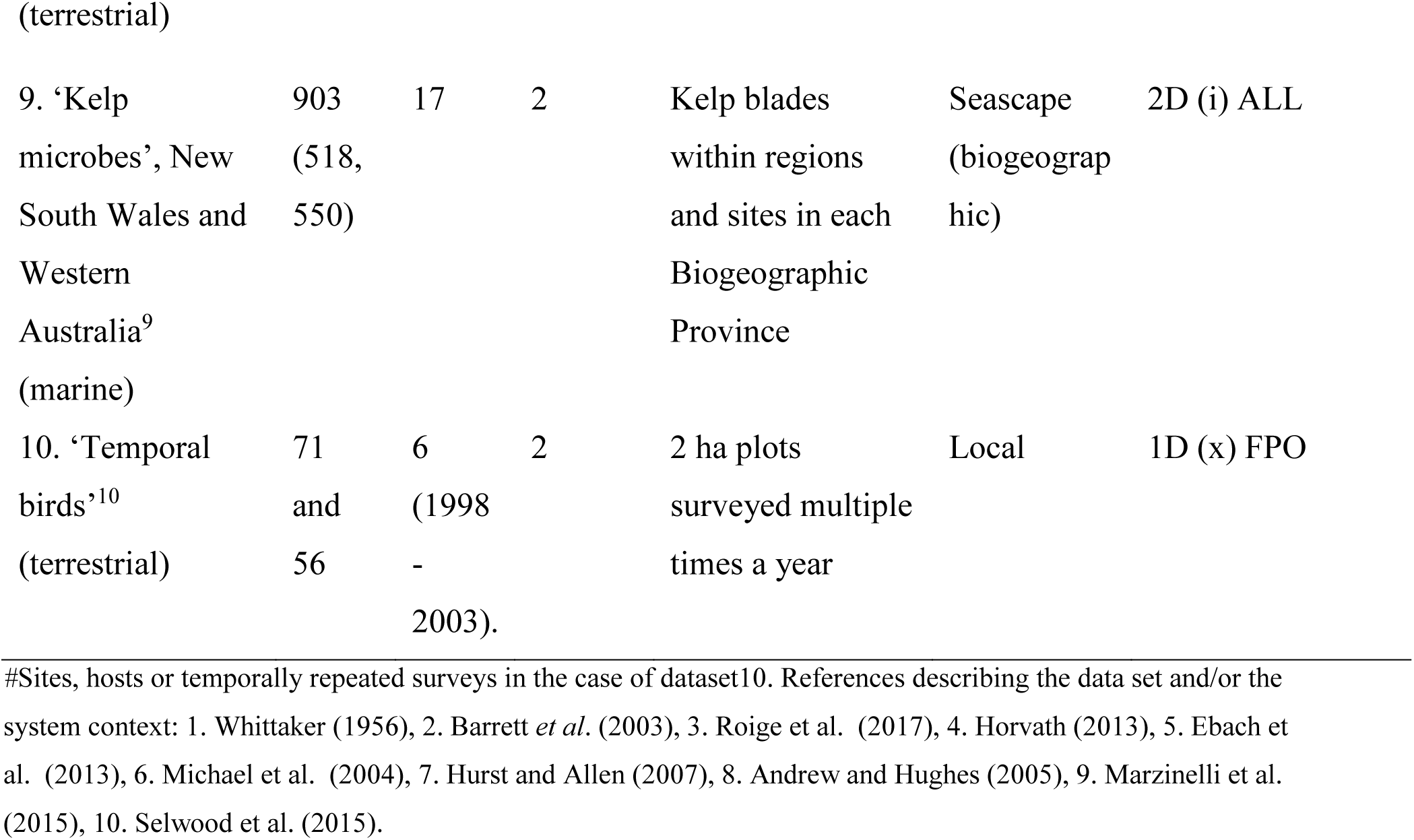
Properties of the ten datasets used to illustrate the application of zeta diversity (in the form of OTU (operational taxonomic unit) by case matrices, see Appendix S2-S5 for further details).

R.H. Whittaker presented the first applications of the concept of beta diversity to quantify turnover in plant communities (which he termed ‘coefficient of community’ now known as Jaccard’s similarity index) in a series of publications spanning the late 1950’s to early 1960’s (Whittaker 1960, 1967). To illustrate the conceptual shift from beta to zeta diversity, we start by using one of the original datasets of Whittaker (1956). Tree community composition was surveyed along an elevational gradient at 122 m intervals at mesic sites in the Great Smoky Mountains, spanning 480 - 1700 m a.s.l. (39 tree species at 11 ‘sites’ or elevational bands) (Table 5 in Whittaker 1956) (referred to from here on as the data set ‘Trees’, see Table 1).

Three different Australian bird datasets were used (Table 1). The first is a selection of atlas data for terrestrial (non-freshwater) species at 25 × 25 km grain, in a 150 km radius around the Sydney Central Business District (33° 51’ 44.4132’’ S, 151° 12’ 31.77’’ E) (Barrett et al. 2003) (‘‘Sydney birds’’, Table 1, Appendix S2). The second dataset uses checklist-type lists of species across the 85 (unequal area) bioregions in the country (Ebach *et al*. 2013) (‘‘Bioregion birds’’, Table 1). The third bird data set includes temporal data for native birds in two catchments in a major river basin in southeastern Australia (‘‘Temporal birds’’, Table 1). These were collected from 2 ha sites over a 6-year period from 1998 to 2003 (Appendix S2), which coincided with a regional drought (Selwood et al. 2015).

Microbial communities (bacterial and archaeal OTUs defined based on a <97% identity of their 16S rRNA genes) associated with the surfaces of common kelp (*Ecklonia radiata*) were examined along the coastline of temperate Australia. Samples from within two marine Biogeographic Provinces (BPs) were examined (alongside the Australian states of New South Wales (NSW) and Western Australia (WA), Appendix S2). Within each BP, 3 regions (spanning ~ 4° latitude or ~ 600km) were sampled with 3 sites per region (Marzinelli et al. 2015) (‘Kelp microbes’, Table 1).

Two very different insect datasets were used. The crop pest data include occurrence records at the level of country, state (province) and island group for over 800 insect pest species of interest to global crop protection (Roige et al. 2017) (‘‘Crop pests’’, Table 1). The second dataset includes insect herbivores (bugs (Hemiptera) and beetles (Coleoptera)) sampled from a single host plant (*Acacia falcata*, data pooled for 120 trees per site) across 12 sites spanning a 1200 km latitudinal extent in Eastern Australia (Andrew and Hughes 2005) (“*Acacia* herbivores*”*, Table 1, Appendix S2).

Plant survey data from Banks Peninsula (New Zealand) includes native and alien plant species (n=1037) from a regular array of plots (n=1338) approximately 1km apart across the extent (~50 × 30 km) of the Peninsula (Wiser et al. 2001) (Appendix S3). (‘‘Plants’’, Table 1).

The ‘Soil metagenome’ data set was generated from twelve, 5 ml soil samples taken as an array within an area of approximately 50 m^2^ in a dry sclerophyll woodland in New South Wales (Australia) (Michael et al. 2004, see for further details on DNA extraction and gene cassette size class screening, assessment and characterisation). The data matrix used is thus based on small, mobile genetic elements (or gene ‘cassettes’) as OTUs versus soil samples (‘‘Soil metagenome’’, Table 1).

Finally, because ecological metrics are increasingly being used for other biological applications (La Salle et al. 2016, Warton and McGeoch 2017), we included a dataset on sub- cellular patterns of turnover that consisted of the presence or absence of DNA hypermethylation (a mechanism used by cells to control gene expression) at nucleotide sites in tissues from patients with and without a metabolic disorder (Table 1). The dataset included the incidence of DNA methylation (‘DNAm’) at CpG (dinucleotide) sites in human occipital cortex tissue from 16 males of a range of ages, with (n=8) or without (n=8) a developmental disorder (autism) (see Ginsberg et al. 2012). Here, age was considered as a relational variable as DNA methylation has been shown to be negatively related to age (Horvath 2013). In this case the OTUs were CpG sites and the tissue from individual patients were the cases (‘‘DNAm’’, Table 1). The question of interest here is – does the distribution of hypermethylation across CpG sites (i.e. compositional turnover) distinguish patients with and without a developmental disorder.

In datasets where a large proportion of the OTUs are shared by the majority of cases (and where the value of zeta would therefore be high at high orders), it may be appropriate to consider this subset of OTUs with a close to saturated distribution as uninformative and to exclude them – as we did for the high proportion of nucleotide sites at which hypermethylation occurred across all patients in the ‘DNAm’ dataset (Appendix S2). These OTUs may otherwise hide the signal in zeta diversity from the whole suite of less common OTUs (see details below). However, in some systems the identification of common suites of species may itself be of interest (Gaston 2010, McGeoch and Latombe 2016). For example, in microbial studies the identification of ‘core microbiomes’ is meaningful (Shade and Handelsman 2012), and wide-ranging components of assemblages are also relevant in invasion biology.

## INTERPRETING ZETA DIVERSITY

### 1 ZETA DECLINE

Zeta decline quantifies how the number of shared OTUs decreases with zeta order, i.e. with increasing number of cases included in the calculation of shared OTUs. Plots of zeta diversity against the order of zeta (i.e. zeta decline) provide information on the form and rate of decline in the average number of OTUs shared across increasing orders of zeta, where orders represent selected pairs (order 2 for value zeta 2), triplets (order 3 for value zeta 3) of cases and so on (Hui and McGeoch 2014).

As a departure point, we used Whittaker’s (1956) tree data to show how traditional pairwise decline using Jaccard similarity compared with the decline in zeta diversity for n-sites, (Fig. 2). [Note that only in this particular and simple case of a one-dimensional data structure and a directional point source sub-sampling scheme, is zeta order (elevational bands in this case) directly comparable to distance along the transect. The data underlying Fig. 2 match the scheme in Fig. 1j, and in this case zeta decline is directly comparable to zeta decay.]. Applying zeta diversity so that it most closely matches the approach used by Whittaker (1967) (Appendix S2,4) revealed a comparatively similar steady decline in species shared beyond the first two elevational bands (Fig. 2). However, normalised zeta across the transect was lower by comparison, as expected given the inclusion of multiple elevational bands in its calculation beyond the second band (normalised ζ_2_ is equivalent to the Jaccard similarity index between the first pair of sites) (Fig. 2). The significance of the difference in interpretation using Jaccard versus zeta diversity is that pairwise comparison of sites underestimates compositional diversity along the elevation gradient. Underestimation of turnover such as this could potentially affect any conservation decision that is made based on relative or comparative levels of heterogeneity, such as the placement of monitoring localities or protected areas (Socolar et al. 2016).

**FIG. 2.**
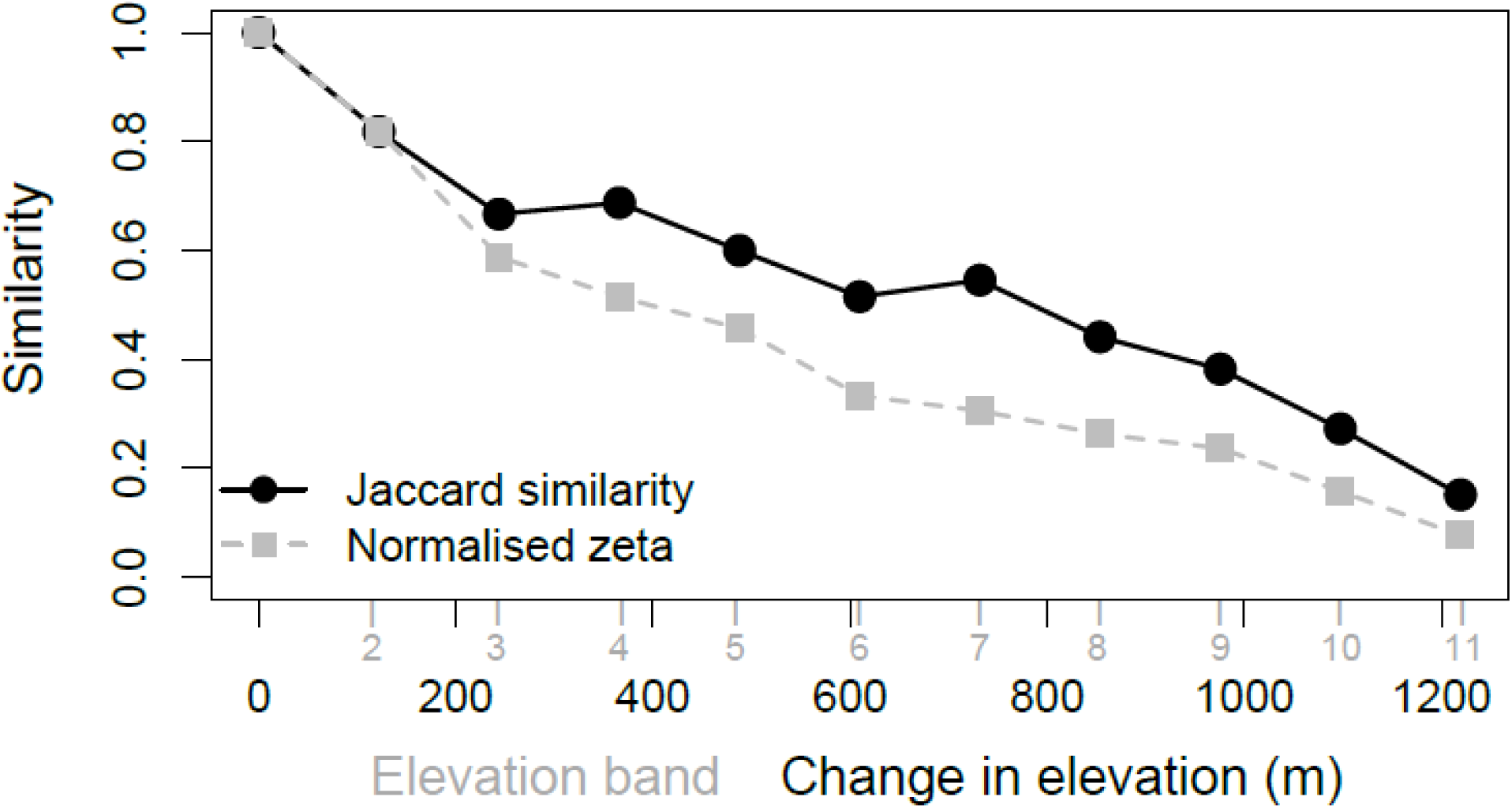
Compositional change in tree species along an elevation gradient in the Great Smoky Mountains, quantified using pair-wise Jaccard similarity as used by Whittaker (1967). Tis is compared with normalised, n-wise zeta diversity decline. Both elevational bands (equivalent to the zeta order in this case) and the distance along the elevational transect (m) can be shown on the x-axis in this case.

In the following sections we examine the ecological interpretation of zeta decline and its parametric form, and introduce the zeta ratio and species retention rate curves built from the zeta ratio.

#### 1.1 The ecological interpretation of zeta decline

Features of interest in zeta decline include: (i) the rate of decline in shared OTUs, particularly across the first few orders, and (ii) if at higher orders the curve reaches or approximates zero or not. The larger the change in the value of zeta across subsequent orders, the greater the relative difference in the numbers of rare versus increasingly common species in the community. At lower orders this provides information on the rate at which rare species are lost from the community. At higher orders, the value of zeta diversity provides information on the existence and size of the common core of OTUs in the community for a particular order, that is of interest itself but also for comparisons within and across datasets.

We used normalised zeta to enable a comparison across datasets (or assemblages with very different richness) with a wide range of OTU richness, including the ‘Crop pest’, ‘DNAm’, ‘Bioregion birds’ and the ‘Soil metagenome’ datasets (Table 1, Fig. 3) (see also Appendix S2,S4). From Fig. 3a, it is apparent that in some cases the average number of OTUs shared across sites declines to zero within the extent of the study system, whereas in datasets with some OTUs present in all sites, zeta converges towards this number of widespread OTUs. The value of zeta at the highest expressed order represents the most common subset of species in the assemblage for that order, i.e. the average number of species shared by large numbers of cases (interpreted as a % using normalised zeta), where large is equivalent to the highest order of zeta expressed in the zeta decline curve.

**FIG. 3.**
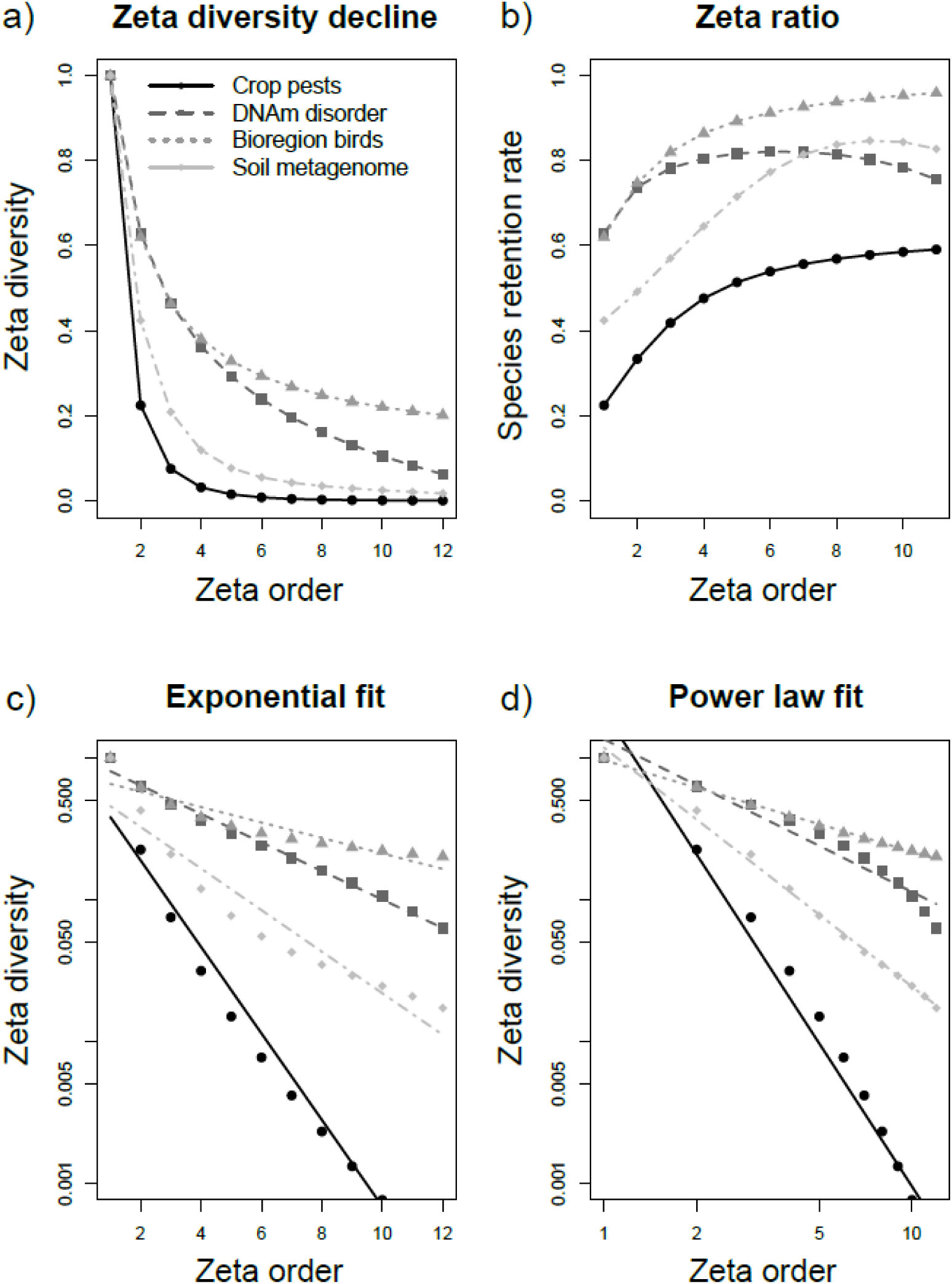
Normalised zeta diversity decline (a) for four data sets (see Table 2) showing how the number of shared OTUs decreases with the zeta order. (b) The species retention rate using the zeta ratio, which shows the degree to which common OTUs are more likely to be retained in additional cases or samples than rare ones with an increase in zeta order. (c,d) The form of decline against exponential (comparatively equal probability of OTUs across cases) or power law fits (comparatively unequal probabilities of the occurrence of OTUs across cases) (shown on log axes using normalised zeta). [Crop pests (circles), DNAm disorder (squares), bioregion birds (triangles), soil metagenome (diamonds)]

The species shared by global crop pest assemblages declined to approximately zero after only 6 orders, and although the rate of decline in the ‘Soil metagenome’ data at a micro scale was somewhat slower, it also declined to approximately zero after ~ 10 orders (Fig. 3a). Ecologically, in both these datasets, the extent of the study exceeds the scale at which communities are structured because the number of shared species declines to zero fairly rapidly. Zeta diversity declined sharply for ‘Crop pests’, with complete turnover in the pest assemblage expected across more than 6 states or countries. Therefore, although there are a small suite of widespread insect crop pests globally shared by several countries, the global composition of pest assemblages actually differs widely (Roige et al. 2017).

By contrast, although zeta decline approximated zero at higher orders of zeta for global ‘Crop pests’, it declined to approximately 20 % of bird species shared by Australian bioregions by order 12 (14% across all bioregions). There was therefore a core set of common bird species (~ 20% or 50 species, across 12 orders) shared across bioregions, shown by the large zeta values for high orders (Fig. 3a, Appendix S5). This long tail of zeta decline for birds represents a set of wide-ranging species that are either habitat generalists (e.g. Australian Owlet-Nightjar (*Aegotheles cristatus*)), or long-range dispersers (e.g. Fairy Martin (*Petrochelidon ariel*)) (Appendix S3). Similarly, but using raw zeta, in the ‘Trees’ data there were a common suite of ~ 5 tree species (Fig. 4a), whereas for ‘Sydney birds’ there were approximately 40 bird species in common on average across combinations of ten or more sites around Sydney (Fig. 4b).

**FIG. 4.**
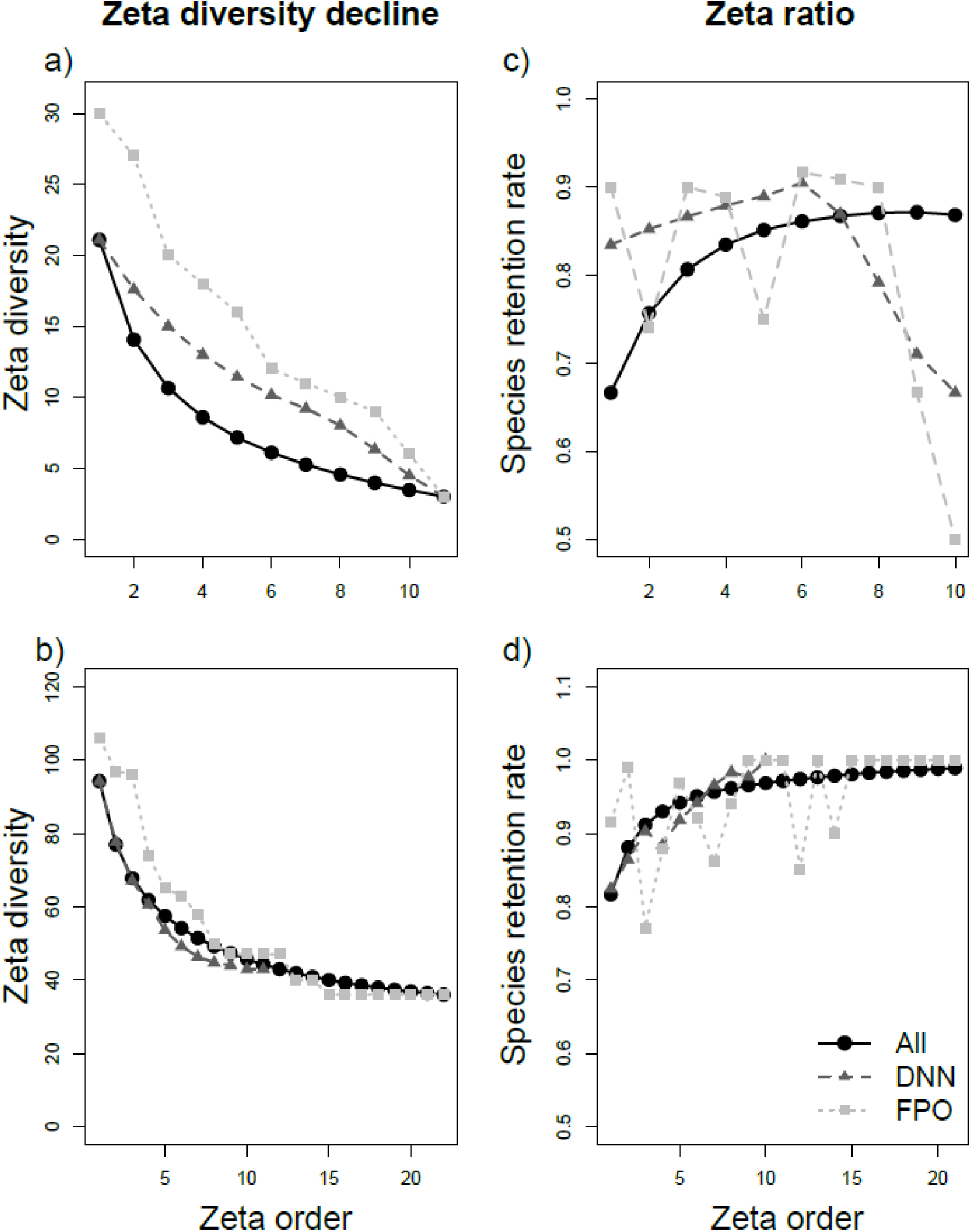
Patterns of compositional change with different data sub-sampling schemes (All, DNN, FPO) are shown for directional data structures (Fig. 1) using zeta diversity decline (a, b) and the zeta ratio plotted as species retention rate (c, d). Data sets used are trees along an elevation transect (a, c; ‘Trees’ Table 1) and bird communities radiating inland from central Sydney (b, d; ‘Sydney birds’ Table 1). Data combination schemes: ALL, all combinations of n sites, DIR, directional nearest neighbour, FPO, fixed point origin (see Fig. 1). The legend in panel d relates to panels a to c.

Intermediate to the other datasets in Fig 3a, shared nucleotide sites at which hypermethylation occurs (‘DNAm’ data) declined more rapidly after zeta order 4 in comparison with bird composition, with fewer than 10% of dinucleotide sites shared by zeta order 12 (Fig. 3a, interpreted further below). Here, the low percentage of shared sites (low zeta diversity) at order 12 is driven in part by the pre-analysis removal of hypermethylation sites shared by all patients, because they are uninformative in the context of this dataset (Appendix S2,S4). This illustrates the importance of biologically-driven decisions on how to treat the data pre-analysis, and the study specificity of how zeta decline is interpreted – at least across this widely divergent set of examples that were used to illustrate the array of possible forms of zeta decline.

#### 1.2 The retention rate based on the zeta ratio

A measure of OTU retention rate can be calculated using the ‘zeta ratio’ (e.g. ζ_2_/ζ_1_). The retention rate curve quantifies the degree to which common OTUs are more likely to be retained across cases than rare ones with an increase in zeta order. Common OTUs are intuitively more likely to be retained in extra samples than rare OTUs, although not necessarily so (dependent to some extent on scale (grain) and species aggregation) (Harte 2008, Hui and McGeoch 2008). By comparing the ratios of zeta diversity values (e.g. ζ_10_/ζ_9_ vs. ζ_2_/ζ_1_), it is therefore possible to assess the extent to which this is the case.

Because the average number of shared OTUs declines with increasing numbers of cases (as in zeta decline), a random species shared by *i* sites has a probability ζ_i+1_/ζ_i_ of still being shared by *i*+1 sites. The zeta ratio plotted against increasing orders is interpreted as the rate (or the probability) at which species are retained in the community as additional cases are included in the comparison. The zeta ratio for a particular order is therefore the probability of retaining (or rediscovering) an OTU of the same order of commonness in additional samples. In addition, as shown in Hui and McGeoch (2014), the specific ratio ζ_1_/ζ_0_ provides an estimate of the probability of discovering new species in additional samples. The abscissa in the species retention rate plot is interpreted slightly differently to the order in zeta decline. For example, the zeta ratio for order nine is interpreted as the probability of retaining species with an occupancy of nine (present at nine sites) in a tenth site, or the probability that these species remain widespread with the addition of another site.

In Fig. 3b, all OTU retention rates start increasing, indicating a rapid loss of rare OTUs and demonstrating that pairwise beta turnover is largely driven by the gain or loss of rare species (consistent with strong modes of rare OTUs, Appendix S5). The probability of retaining common species is much lower for ‘Crop pests’ than ‘Bird bioregions’, but the rates of common species retention for both these datasets start to asymptote beyond order 6 (Fig. 3b). The retention rates for the ‘Soil metagenome’ and ‘DNAm’ data increase and then start to decline (i.e. show signs of becoming modal, for a stronger example of this form of species retention curve see Fig. 5b, beetles). This means that at higher orders there is a decline in the probability of retaining common species in the community with an increase in order (or a decrease in the rate of OTU retention) (Fig. 3b). Across all the datasets examined (see also examples in Fig. 5), three general forms of retention rate curves were observed, (i) increasing (e.g. the bugs in the *‘Acacia* herbivores*’* data, Fig. 5b), (ii) asymptotic (e.g. ‘Bioregion birds’ and ‘Crop pests’, Fig. 3b) and (iii) modal (e.g. beetles in the *‘Acacia* herbivores*’* data and to a lesser extent the ‘DNAm’ and ‘Soil metagenome’ data, Figs 3b, 5b).

**FIGURE. 5.**
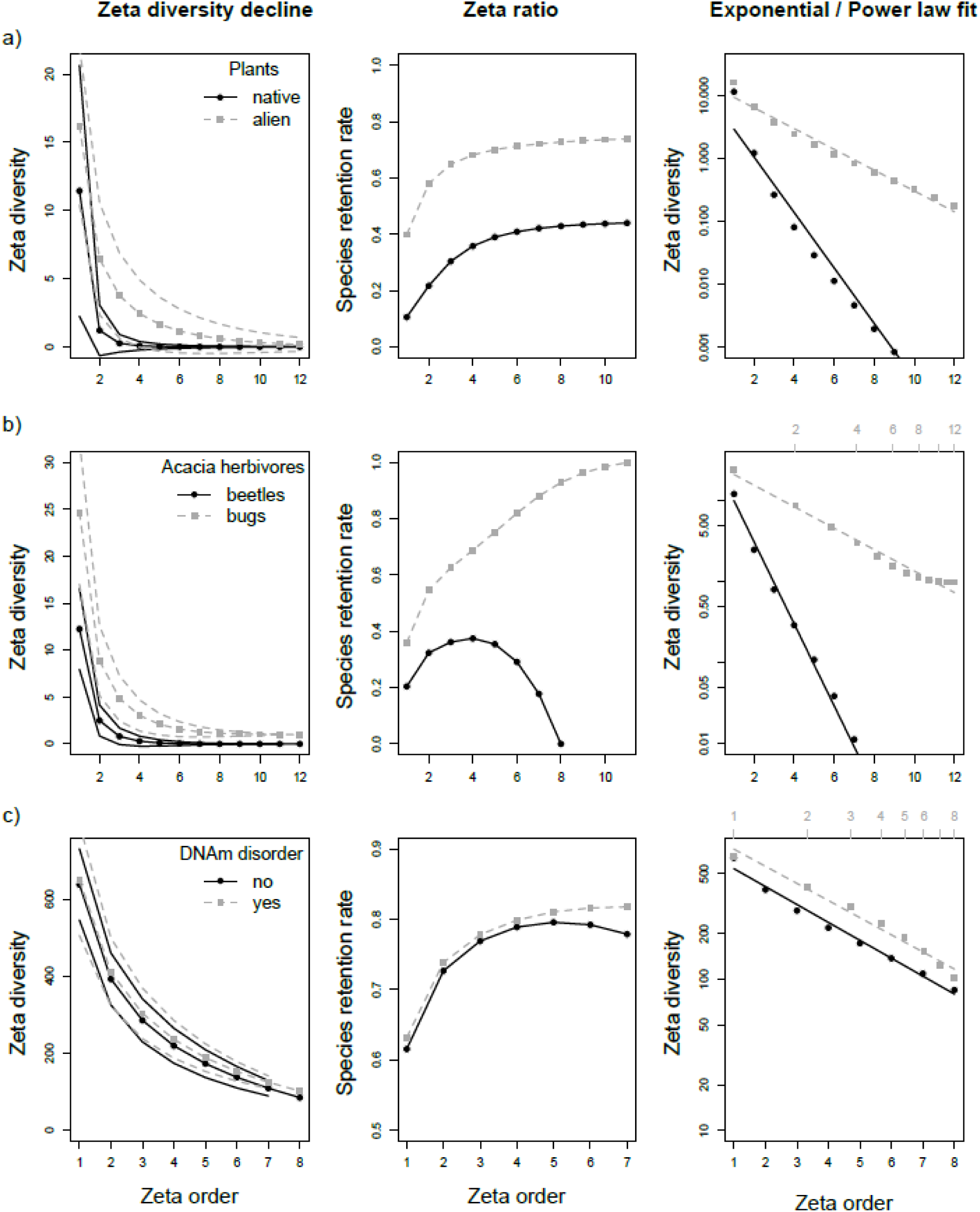
Comparisons of zeta diversity decline between OTU or case groups in three data sets, along with species retention rate using the zeta ratio, and exponential versus power law fit (on log axes): a. alien and native plants on Banks Peninsula; (b) *Acacia* herbivores (beetles and bugs) across a latitudinal gradient, and (c) DNA hypermethylation sites in patients with and without a disorder. The data sub-sampling scheme in all cases is ‘ALL combinations’ (Fig. 1i).

Within a study system the three types of retention rate are likely to be a continuum, shifting from increasing to asymptotic if a core set of common OTUs remain for a particular zeta order, and either directly from increasing to decreasing, or via a modal curve, when moving beyond the footprint of the most common suite of OTUs in the community. The biological significance of these will be study dependent, but can generally be interpreted as follows: An increasing curve indicates that common OTUs are more likely to be retained in additional samples than rare ones, and as a result perhaps that the sampling extent is narrower than the metacommunity, or that site selection is relatively homogenous and well characterised by habitat specialists (Myers and LaManna 2016). In an asymptotic curve, an asymptote of 1 indicates the presence of common species over all sites, whereas an asymptote < 1 indicates that common and intermediate species are equally likely to be retained in subsequent sites or samples. A modal curve indicates that for high orders of zeta, the most common OTUs are less likely to be retained when adding sites, i.e. the study extent encompasses the community or metacommunity (Appendix S3).

Examining plots of the zeta ratio expressed as species retention rate curves is particularly useful for visualising turnover at higher orders (which in zeta decline curves may be difficult to distinguish because the values of zeta are low) and highlights potential differences between the zeta declines of related datasets. This is apparent in the ‘*Acacia* herbivore’ (beetle) and the DNAm data (Fig. 5b,c), which revealed patterns at higher zeta orders that were not apparent from the zeta decline curves.

#### 1.3 Effect of data subsampling scheme on zeta decline and retention rate

As outlined above, when applying zeta diversity it is important not only to use an appropriate survey design (as for any ecological study), but also to consider the appropriate data subsampling scheme for the system and question of interest (Fig. 1). Comparing zeta diversity decline using three data sub-sampling schemes on the ‘Trees’ and ‘Sydney birds’ data (Fig. 4) illustrates the shallower rate of decline over all combinations (ALL) and using nearest neighbours (DIR), than using the fixed point origin (FPO). This is a consequence of spatial clustering of species and the continuity of ranges, particularly of the more common species across the transect. This is apparent for both the one-dimensional ‘Trees’ data (Fig. 4a), and the two-dimensional data structure for ‘Sydney birds’ (Fig. 4b).

Comparing the results from three subsampling schemes on the ‘Trees’ dataset illustrates the potential ecological value of retention rate curves (Fig. 4a,c). The zeta retention rate curve is particularly striking, with a rapid decline in the rate of species retention beyond zeta orders 6-9 for the DIR and FPO schemes (Fig. 1i, j). This is not apparent from the zeta decline curve (Fig. 3a) nor from the zeta ratio using the ALL scheme (Fig. 4c). For comparison, the zeta ratio identifies no sudden shift in bird composition in Fig. 4d for any of the three subsampling schemes. The rate of species retention stabilises beyond zeta order 10, demonstrating the absence of any conspicuous ecotone or dispersal barrier across the urban area encompassed by these bird data.

Whittaker (1967) concluded from his analysis of the change in Jaccard similarity in tree composition (from the lower elevational origin) across the Smoky Mountains elevational transect, that there was broad overlap in species distributions along the gradient. He remarked on the ‘striking’ straight-line relationship between log similarity and the elevational gradient. However, although Whitaker (1967) interpreted the patterns of Jaccard-based distance decay (as shown in Fig. 2) as the existence of ‘broadly overlapping’ species distributions across the transect, he also anecdotally pointed out the existence of a switch in dominance from cove forest species to gray beech and a suite of small tree species at ~ 1400 m a.s.l. along the transect (Whittaker 1960). This coincides with the abrupt shift in species composition between 976 m - 1098 m detected by the zeta ratio and shown by the sharp decline in species retention rate for the DIR and FPO subsampling schemes (Fig. 4c). In the ‘Trees’ data, the retention rate of zeta diversity computed with the appropriate subsampling scheme thus enabled the identification of the ecotone noted by Whittaker (1960), by better capturing the contribution of common species to turnover along the gradient, in comparison with pairwise beta diversity (equivalent to ζ_2_).

Spatially or environmentally structured sampling schemes affect the form of both the zeta decline and the retention rate. These may therefore be compared with the ALL sub-sampling scheme to test mechanistic explanations of turnover (McGill and Nekola 2010, Myers and Manna 2016, Latombe et al. 2017).

#### 1.4 The parametric form of zeta decline

The parametric form of zeta decline as best fit by either a power law or exponential relationship provides insight on the relative probability of OTU (species) occurrences across cases (sites), and may be used to test hypotheses about the extent to which biological matrices or communities are structured (Hui and McGeoch 2014). Power law and exponential parametric forms have been shown to most often best fit decline curves, although other distributions are possible (Hui and McGeoch 2014). Estimated using ALL site combinations (Fig. 1a, g), the parametric form of decline is interpreted as OTUs having the same (exponential) or unequal (power law) probability of being observed across cases.

The ‘DNAm’ data were better fit by an exponential than power law (AIC -39.77 versus - 18.93), whereas the difference was marginal for ‘Crop pests’ (AIC -1.96 exponential versus - 1.47 power law) (Fig. 3c,d). This result shows, at least for the ‘DNAm’ data, a lack of structure in the matrix and that there are approximately equal probabilities of hypermethylation occurring at any nucleotide site. The two other datasets were better fit by a power law (AIC value differences > 30) (Fig. 3c,d), demonstrating some structure in the ‘Bioregion birds’ and ‘Soil metagenome’ datasets and uneven probabilities in the occurrence of OTUs across cases.

Comparatively equal probabilities of the occurrence of species across sites (exponential form) has been suggested to be associated with stochastic assembly processes, whereas habitat heterogeneity and niche differentiation processes are more likely to produce a power law form of zeta decline in natural communities (Hui and McGeoch 2014, for comparable mechanistic beta-diversity based interpretations see Munoz et al. 2008, Nekola and McGill 2014). The fit can also be used to test the scale dependence of OTU incidence in the community; exponential reflects scale independence of species retention, whereas the power law reflects non-independence across cases, and an increasing probability of retaining more common OTUs at finer scales (Hui and McGeoch 2008, McGlinn and Hurlbert 2012). The relationship between the parametric form of zeta decline and mechanistic process in biological systems requires further testing. As with any inference of process from pattern in ecology, clear hypothetical frameworks and strong inference approaches should be used to support the interpretation of the parametric form of zeta decline in this way.

#### 1.5 Within-system comparisons of zeta decline

In the previous examples (Fig. 3) we contrasted datasets that would not normally be included in the same study, to illustrate the range of possibly forms of zeta diversity decline and retention rate. Here, using raw rather than normalised zeta, we use three examples to compare zeta diversity within individual datasets across different OTU (Fig. 5a, b) or case (Fig. 5c) groups (using ALL combinations (Fig. 1i)).

##### Example 1. An invaded plant community

Clear differences are apparent in compositional change between alien and native ‘Plants’ (95% CI = [1.74, 1.95] in a linear model, Fig. 5a, Appendix S4). Alien turnover declines more slowly than native plant composition. Here, although there are over half as many alien as native plant species on Banks Peninsula (Wilson 2009), there were higher values of zeta diversity (more alien species in common than natives) and slower turnover in alien compared to native plant species composition. Alien turnover declines more slowly than native plant composition, and the zeta ratio shows that within the alien plant subset, common species are more likely to be retained across sites (by between ~40-70%) than in the native plant subset (~10-40%) (Fig. 5a). Both native (ΔAIC = 3.96) and alien (ΔAIC = 2.42) zeta decline are better fit by an exponential than power law, suggesting little structure in the plant community at the scale of this study, i.e. species on average have comparatively equal (albeit low) probabilities of being found across sites (Fig. 5a).

##### *Example 2. Insect herbivores on* Acacia

Clear differences are apparent in compositional change between the two groups of ‘*Acacia* herbivores’ (95% CI = [1.81, 1.94] in a linear model, Fig. 5b, Appendix S4). For *‘Acacia* herbivores’, the decline in beetle species shared across the gradient is very rapid (exponential, ΔAIC = 20.01), reaching a zeta diversity of zero by order 10, in contrast to slower decline in compositional similarity in bugs across the same gradient (power law, ΔAIC = 26.49) (Fig. 5b). Whereas the species retention rate in bugs is increasing, for beetles the retention rate drops beyond zeta order 5 (Fig. 5b). The probability of retaining beetle species in the assemblage (zeta ratio) beyond order 4 declines rapidly, suggesting complete turnover in the composition of beetles on *Acacia* within the extent of this study (Fig. 5b). Low prevalence and abundance of beetles in samples (Andrew and Hughes 2005) is a plausible explanation for the strong decline in species retention and lack of structure (i.e. exponential zeta decline) observed in these data.

##### Example 3. Hypermethylation at nucleotide sites

There was little difference in compositional turnover of hypermethylation sites across patients with (parametric form not distinguishable, ΔAIC= 0.01) and without (exponential, ΔAIC = 3.99) a metabolic disorder evident from a comparison of their zeta decline and retention rate curves (Fig. 5c). Using disorder status (binary) and patient age as predictors for zeta order 2 to 4, status was not significant (supporting the multivariate analysis-based findings of Ginsberg et al. 2012), whereas age was a significant predictor of ζ_3_ (95% CI = [-66.07, -25.79]) and ζ_4_ (95% CI = [-58.19, -25.81]), but not ζ_2_ (95% CI = [-98.44, 8.47]). The general prevalence of a relationship between DNA methylation and age is well known (Horvath 2013), but was detected here only for orders of zeta greater than 2, i.e. not detected by beta diversity (ζ_2_). This demonstrates that examining the full spectrum of rare to intermediate and common OTUs as they contribute to driving compositional change is more information rich than quantifying pairwise compositional turnover alone.

### 2 ZETA DECAY

Zeta decay quantifies change in the number of OTUs shared with increasing distance between sites (or time between surveys) for different orders of zeta. Zeta decay is conceptually similar to distance decay (Nekola and McGill 2014), or species–time relationships and time decay (Shade et al. 2013), and provides information on the spatial or temporal extent of communities. It also provides information that can be used to design the spatial and temporal dimensions of sampling schemes to capture features of biodiversity change of interest. Zeta decay, or a plot of zeta diversity across sets of cases that are different distances or times apart, is represented with each zeta order as a different decay curve. In temporal decay the curves represent the change in number of shared OTUs across subsequent surveys or time periods (this can vary with sampling scheme, see Fig. 1). Note that the ends of zeta decay curves, in particular the longer distance end, are usually associated with greater uncertainty because there are comparatively fewer cases this maximum distance apart than there are combinations of cases shorter distances apart (the same problem of unequal power across classes occurs in estimates of autocorrelation series, Legendre 1993).

For orders i > 2, the distances between pairs of *n* sites are combined using, for example, mean distance (other options are the extent of occurrence (EoO) by the cases under consideration, or the maximum distance of cases apart). This must be considered when interpreting the effect of distance on zeta diversity as the order increases (Latombe et al. 2017).

Using zeta diversity decay, spatial and temporal compositional similarity for each order of zeta illustrates differences in the form of decay for the rare to more widespread OTUs in the community over time or distance (Fig. 6). Characteristics of interest are (i) the shape and rate of change (slope) of decay, and how this differs across orders of zeta, (ii) the absolute distance (or time) over which this decay in the similarity of OTU composition occurs, and (iii) the presence or absence of periodicity in the curves.

**FIG. 6.**
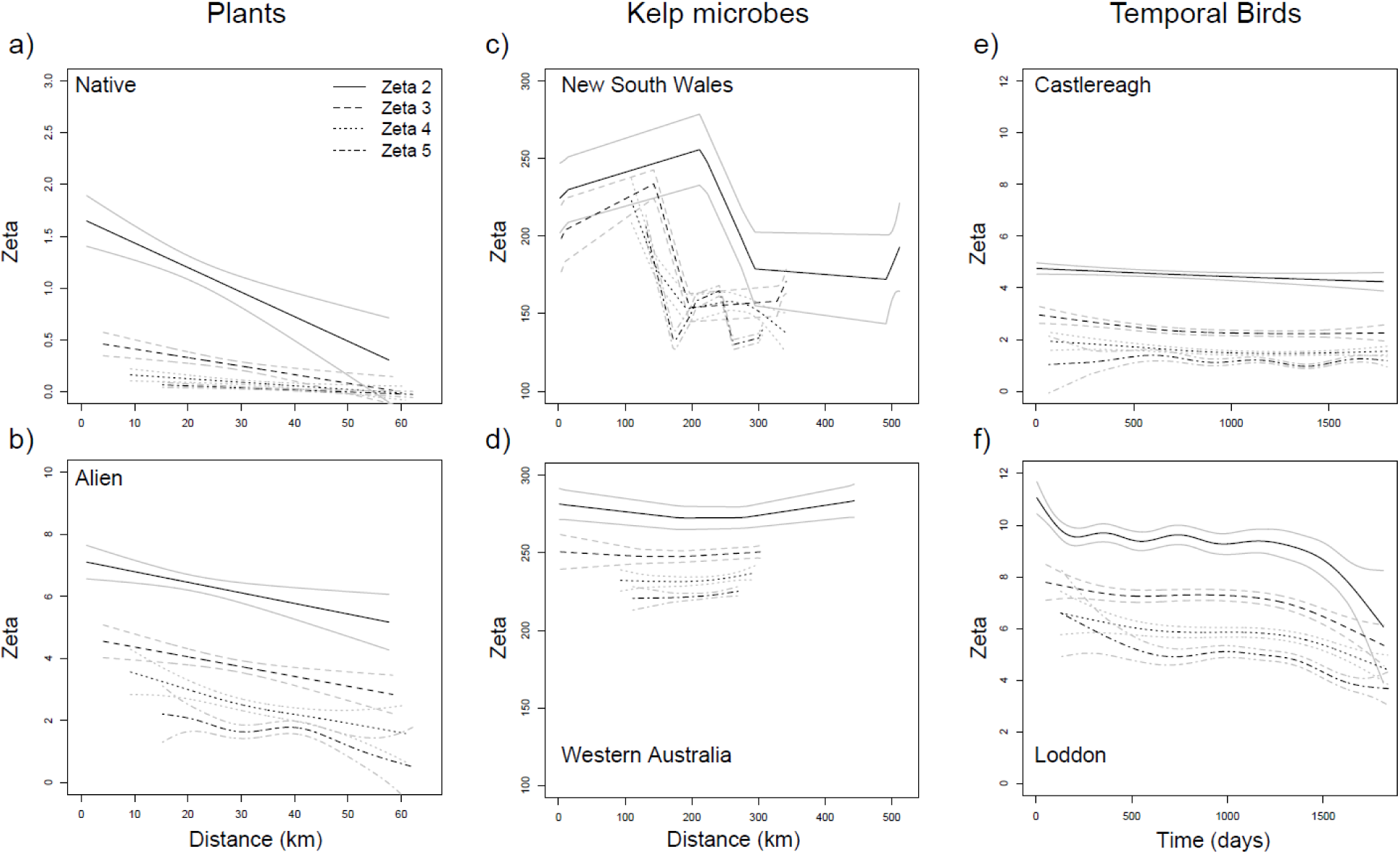
Zeta diversity decay over space and time, for zeta orders 2 to 5, showing change in number of OTUs shared with increasing distance between sites (or time between surveys). (a-b) Alien and native plant species on Banks Peninsula (New Zealand); (c,d) microbial communities associated with kelp in two Australian marine biogeographic regions (New South Wales (east) and Western Australia (west)) using ALL combinations (see Fig. 1i); (e,f) temporal decay in bird communities in two catchments (Castlereagh River, 5% below average rainfall; Loddon River, 10% below average rainfall) over the course of a regional drought (1998-2003) (turnover relative to first year of the drought, i.e. fixed point origin scheme FPO, Fig. 1j)). Note that using mean distance for higher orders (*i* > 2) of zeta (c,d) results in the increasingly narrow decay curve with increasing distance or time (see text).

#### 2.1 Patterns of zeta decay

Four general patterns of zeta diversity decay were apparent in the examples used (Fig. 6, Appendix S4). First, decay was shallow to absent in Fig. 6d,e across zeta orders 2 to 5. Second, in Fig. 6a,b decay was evident and monotonic for zeta 2 and to a lesser extent for zeta orders 3-5. Third, decay was markedly periodic in Figs 6c and 6f. Finally, differences in the average value of zeta across zeta orders 3-5 ranged from comparatively large (e.g. Fig. 6b) to small (e.g. Fig. 6a). These patterns are interpreted in the context of their datasets below.

The patterns of distance decay for alien and native ‘Plants’ (Fig. 6a,b) confirm the interpretation of zeta decline for this data set in Fig. 5, i.e. more shallow compositional turnover in aliens than natives. Here, however, the difference in rates of decline are calibrated against distance, enabling scale-specific comparisons of distance decay across species groups. Over distances of 20 km, on average there are 2 - 6 alien species shared (across zeta orders), whereas there are fewer than 1 to just over one native species shared by sites this far apart (Fig. 6a,b). The relative distances in zeta values across orders 3-5, especially at larger distances in Fig. 6b, illustrate that there are more ubiquitous species (both locally and regionally widespread) in the alien than the native community. If on-ground surveys were to extend beyond the current sample extent, one might expect therefore to discover new rare species at a faster rate than new alien species (with the assumption that local species richness remains similar in the newly surveyed sites). These difference in decay slope between native (steep) and alien (shallow) ‘Plants’ is in the direction that one might expect given the tendency for alien and invasive plant species to have broader niches and geographic ranges (Lockwood et al. 2005).

Patterns of distance decay for ‘kelp microbes’ differed markedly at the scale examined across the eastern and western bioregions of Australia (Fig. 6c,d). The steep decline in average numbers of shared OTUs (both rare to more widespread, i.e. from zeta order 2 upwards) over distances of 150-300 km along the coast of NSW suggests marked patchiness in community structure at this scale. By contrast, the rate of distance decay in the WA community was shallow and consistent across the different orders, in spite of high total and average OTU richness in the region (Fig. 6d). On average, the number of shared OTUs was higher and more consistent with distance in Western Australia (total richness 550 OTUs, mean±s.d. = 346.88±23.49) compared with New South Wales (518 OTUs, 288.33±60.02). Compositional change in higher orders of zeta tended to mimic decay in ζ_2_, although over a more narrow range of distances as a consequence of plotting decay against the mean distance across the *i* samples (Fig. 6c,d). Curves with a clear shift or periodicity (where the width of the error intervals should broadly not exceed the amplitude of the shift or period) suggest the presence of a dispersal barrier, a shift in environmental conditions, patchiness or temporal periodicity of some form (Nekola and White 1999). For example, the striking difference between decay curves for kelp microbes between NSW and WA can be explained by distinctly different current systems between the coasts that drive the dispersion of kelp microbes in different ways (Thompson et al. 2011) (differences in the relative distances across sites may also play a role, Appendix S6).

Although the average number (±s.d.) of bird species shared over time (‘Temporal birds’) was similar at the two catchments in the drought-affected river basin in Australia (12.18±3.31 at Castlereagh, 14.81±3.14 at Loddon), compositional similarity was lower (i.e. fewer shared species across years) at Castlereagh than at Loddon River (Fig. 6e, f). Turnover in assemblage composition was comparatively stable over the course of the drought at Castlereagh (shallow decline in zeta diversity), whereas the temporal decay in similarity was more marked at Loddon, particularly in the first year of the drought (1998-1999, over the first ~ 356 days, Fig. 6f, Appendix S6). After ~3.5 years at Loddon, the average number of species in common with the assemblage at the start of the drought started to decline again (this is particularly apparent for zeta orders 3-5). Periodicity in the zeta decay of the more drought affected Loddon bird community suggests some resistance after an initial perturbation during the early stages of the drought (see Selwood et al. 2015), with higher turnover (fewer shared species) over time further into the drought period. The drought was not as severe at Castlereagh, and here the bird community appeared to be comparatively resistant, with very little temporal decay (Selwood et al. 2015).

The differences in zeta decay across zeta orders in these examples illustrates the relative differences in the contributions of rare to more common OTUs to turnover with distance and time. The examples revealed shallow to steep decay slopes, as well as monotonic versus periodic patterns of decay. Although here we speculate on what may be driving the patterns found, drivers of patterns in zeta diversity decay can be formally tested using multi-site generalised dissimilarity modelling, a form of direct gradient analysis, in which zeta diversity is regressed against environmental differences and distance (Latombe et al. 2017). Direct gradient analysis on species composition is traditionally performed using Redundancy Analysis or Canonical Correspondence Analysis (Legendre and Legendre 2008), and relies on linear regression approaches. More recently, Ferrier et al. (2007) proposed a flexible, non-linear version of direct gradient analysis named Generalised Dissimilarity Modelling (GDM). GDM predicts pairwise beta diversity (e.g. Bray-Curtis Dissimilarity) from environmental difference between sites, while accounting for the impact of the environmental gradient on the effect of the environmental difference on compositional turnover. However, since this approach relies on pairwise comparisons of sites, the outputs remain mainly driven by rare species. Extending GDM to zeta diversity to create Multi-Site Generalised Dissimilarity Modelling (MS-GDM, Latombe et al. 2017) enables the identification of differences in the abiotic variables structuring compositional change in rare to common OTUs. Being able to disentangle spatial and temporal trends in rare to common species has significant potential value, given the important role of common species in delivering ecosystem services (McGeoch and Latombe 2016).

## CONCLUSION

When a new approach is proposed that for the first time quantifies, or quantifies differently, a component of biodiversity, the outcome of its application to a range of biological or ecological scenarios becomes of interest, because of the potential that it may reveal new insights about biodiversity. Here we have shown using a diverse range of empirical examples that zeta decline, the zeta ratio and retention rate, the parametric form of zeta decline and zeta decay provide a range of insights on the nature of continuous compositional turnover and the scaling of biodiversity structure. We have also shown how its application reveals patterns of turnover that are not apparent using measures of compositional change for a fixed number of, usually pairwise, cases. The broad range of applications and insights that can be derived using zeta diversity on any incidence matrix will, we hope, also contribute to further development of general theory on the scaling of biotic heterogeneity.

In spite of substantial focus on biodiversity change over the recent period (Butchart et al. 2010), trends in spatial and temporal turnover across scales, from local to global, remain poorly supported by empirical studies (Dornelas et al. 2013, McGill et al. 2015). Our intention here was to show how zeta diversity can contribute to filling this gap when used to study trends in turnover across multiple cases and levels of biological organisation. Along with insights provided by decomposing compositional change into richness and replacement components (e.g. Baselga 2010, 2013), future progress in modelling and hypothesis testing using zeta diversity will be made using combinations of empirical and simulation modelling. With the growing interest in biodiversity turnover and the importance of common species in an increasingly homogenised world (McGeoch and Latombe 2016), advances in ways to measure compositional change and the dynamics of common species, such as zeta diversity, are timely.

## ACKNOWLEDGEMENTS

We thank David Warton, Mark Westoby, Marie Henriksen, Steven Chown and David Baker for comments on the work. M.M., G.L. and C.H. acknowledge support from the Australian Research Council’s *Discovery Projects* funding scheme (project number DP150103017), N.R.A. from the Australian Research Council’s *Discovery Projects* funding scheme (project number DP160101561), and C.H. from the National Research Foundation of South Africa (no. 89967 and 76912). S.N. is supported by a Future Fellowship (FT130100268). M.R. received financial support from the European Commission (Erasmus Mundus partnership NESSIE, ref. 372353-1- 2012-1-FR-ERA MUNDUS-EMA22) and BPRC travel grant 2014. Refer to Appendix S2 for comprehensive data use acknowledgements.

## Footnotes

1 We use the term turnover in its broadest sense to mean change in composition of elements across sites or over time, including both richness dependent and independent components

## SUPPORTING INFORMATION

Additional Supporting Information may be found.

APPENDIX S1. Main pathways for the use of zeta diversity, from the consideration of data structure, to the sub-sampling scheme for combining data to calculate zeta, and how it may be expressed and interpreted.

APPENDIX S2. Sources and accessibility of the ten datasets used, as well as data treatment details for the purpose here of applying zeta diversity.

APPENDIX S3. Further detail about each dataset and the specific zeta diversity analyses applied to each.

APPENDIX S4. Spatial and temporal distribution of cases across datasets.

APPENDIX S5. Zeta decline and associated zeta ratio and species retention rates for all datasets, in each case including the maximum number of zeta orders possible based on the number of cases in the dataset.

APPENDIX S6. Occupancy frequency distributions for each dataset and subset used in analysis.

